# Responses of ground-dwelling spider assemblages to changes in vegetation from wet oligotrophic habitats of Western France

**DOI:** 10.1101/380212

**Authors:** Denis Lafage, El Aziz Djoudi, Gwenhaël Perrin, Sébastien Gallet, Julien Pétillon

## Abstract

While many arthropod species are known to depend, directly or indirectly, on certain plant species or communities, it remains unclear to what extent vegetation shapes assemblages of other taxa, notably spiders. In this study, we tested whether the activity-density, composition, and diversity of ground-dwelling spiders were driven by changes in vegetation structure. Field sampling was conducted using pitfall traps in bogs, heathlands, and grasslands of Brittany (Western France) in 2013. A total of 8576 spider individuals were identified up to the species level (for a total of 141 species), as well as all plant species in more than 300 phytosociological relevés. A generalised linear model showed that spider activity–density was negatively influenced by mean vegetation height and mean Ellenberg value for moisture. Indices of diversity (α, β, and functional diversities) increased with increasing vegetation height and shrub cover (GLMs). Variables driving spider composition were mean vegetation height, dwarf shrub cover, and low shrub cover (results from a redundancy analysis). Spiders, some of the most abundant arthropod predators, are thus strongly influenced by vegetation structure, including ground-dwelling species. Although later successional states are usually seen as detrimental to local biodiversity in Europe, our results suggest that allowing controlled development of the shrub layer could have a positive impact on the diversity of ground-dwelling spiders.

## Introduction

Globally, increases in plant species diversity or structural heterogeneity are often correlated with an increase in species richness of animals (Southwood et al. 1979; Madden and Fox 1997). The architectural or structural heterogeneity of plants, which is likely correlated with both plant-species diversity and productivity (Lawton 1983), can be an important determinant of arthropod diversity and abundance at different trophic levels (Lawton 1983). Many arthropod species depend, directly or not, on vegetation, and it consequently shapes their assemblages (Lewinsohn et al. 2005). This is especially obvious for phytophagous taxa, but has also been shown for other groups using vegetation as shelter or, in the case of spiders, for building their webs. Spider assemblages of vegetation-dwelling and web-building guilds are known to be shaped by vegetation structure, but ground-dwelling spiders are ideal models to test whether vegetation also drives community structure and composition of ground-active predators. While strong relationships have been reported previously between web-building spiders and vegetation (e.g. Ávila et al. 2017), other studies reported a weak effect of vegetation structure on spider diversity (Rodrigues et al. 2014) and no bottom-up effect of vegetation biomass on spiders (Lassau and Hochuli 2008; Lafage et al. 2014; Sousa-Souto et al. 2014). Ground-dwelling spiders are known to react to several local, abiotic factors, such as pH, disturbance, soil structure or moisture level (Schaefer 1990; Andersen 1995; Paquin and Coderre 1997; Pétillon et al. 2008). Their abundance and species richness also respond to the depth and complexity of the litter layer (Uetz 1976, 1979; Hurd and Fagan 1992) which are often related to vegetation complexity. For instance, Langellotto and Denno (2004) found a positive relationship between the abundance of hunting spiders and vegetation heterogeneity. Blaum et al. (2009) also found a hump shape relationship between both spider abundance and species richness and shrub cover.

Numerous studies have tried to understand the determinants of assemblages’ composition and local species richness, i.e. α-diversity (e.g. Hendrickx et al. 2007, Jiménez-Valverde et al. 2010, Pétillon et al 2008). There are fewer studies dealing with β-diversity and functional diversity (McKnight et al. 2007), but their number has increased in recent years (e.g. Hendrickx et al. 2007; Boieiro et al. 2013; Braaker et al. 2013; Woodcock et al. 2013; Lafage et al. 2015). In this study, we tested whether assemblages of ground-dwelling spiders are shaped by changes in vegetation structure along a successional gradient. We took advantage of the monitoring systems of heathlands and grasslands (i.e. fairly stable habitats where landscape factors are likely less determinant than local factors for spiders: Horváth et al. 2015) in natural reserves to test whether a set of variables derived from the surrounding vegetation was able to predict changes in spider structure and composition at ground level. We therefore chose to investigate assemblage composition, α-diversity, β-diversity, and functional diversity activity-density. Spider activity-density was expected to be positively influenced by vegetation complexity due to a higher abundance of prey. Species composition and α-diversity were expected to be positively influenced by local abiotic factors. Indeed, α-diversity describes within-habitat diversity (MacArthur 1965) and is mainly driven by local processes while β-diversity is generally thought to be driven by both local and landscape factors, the latter being the predominant factor for spiders and carabids (Lafage et al. 2015). We expected a weak or null link between vegetation structure and β-diversity. Finally, functional diversity was expected to be positively linked to vegetation complexity as this would allow more guilds to coexist (Cardoso et al. 2011).

## Material and Methods

### Study sites and habitats

Samples were taken in three Special Areas of Conservation (SAC) in the inner part of Brittany (Western France) at the head of drainage basins, including two natural reserves: the bogs of Langazel (LG) and the fens and heathlands of Lann Bern and Magoar-Penn Vern (LB) (Fig. 1). Both sites comprise colluvial and peaty plains crossed by streams and alluvial ‘streaks’. They are composed of wet oligotrophic habitats including large areas of *Ulicion minoris* heathlands (EUR 28 4020) and *Juncion acutiflori* rush pastures (EUR 28 6410) sometimes in a mosaic with small patches of blanket bog communities (*Oxycocco palustris-Ericion tetralicis*).

**Figure 1.**
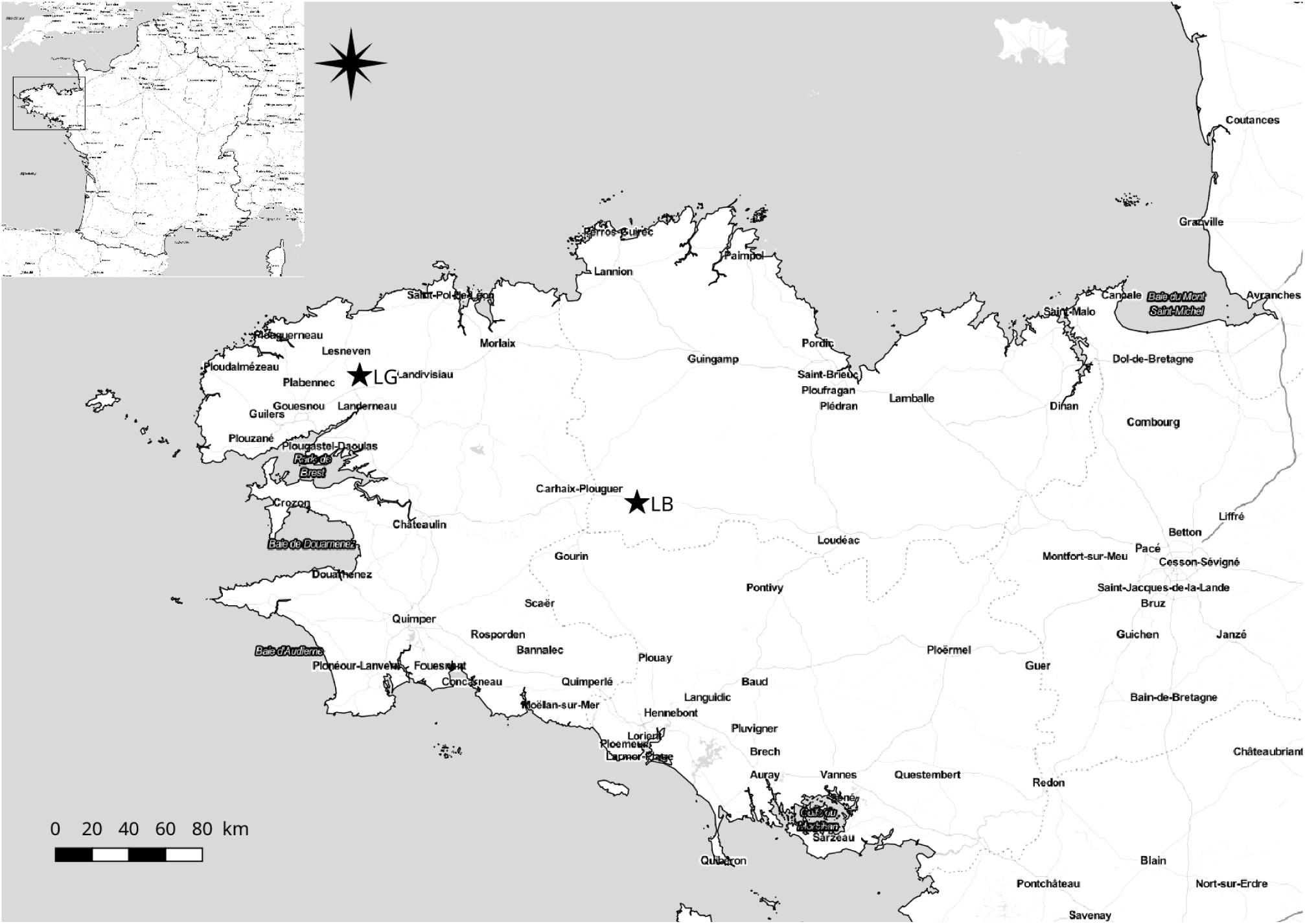
Site locations. LG: bog of Langazel. LB: fens and heathlands of Lann Bern and Magoar-Penn Vern.

**Figure 2.**
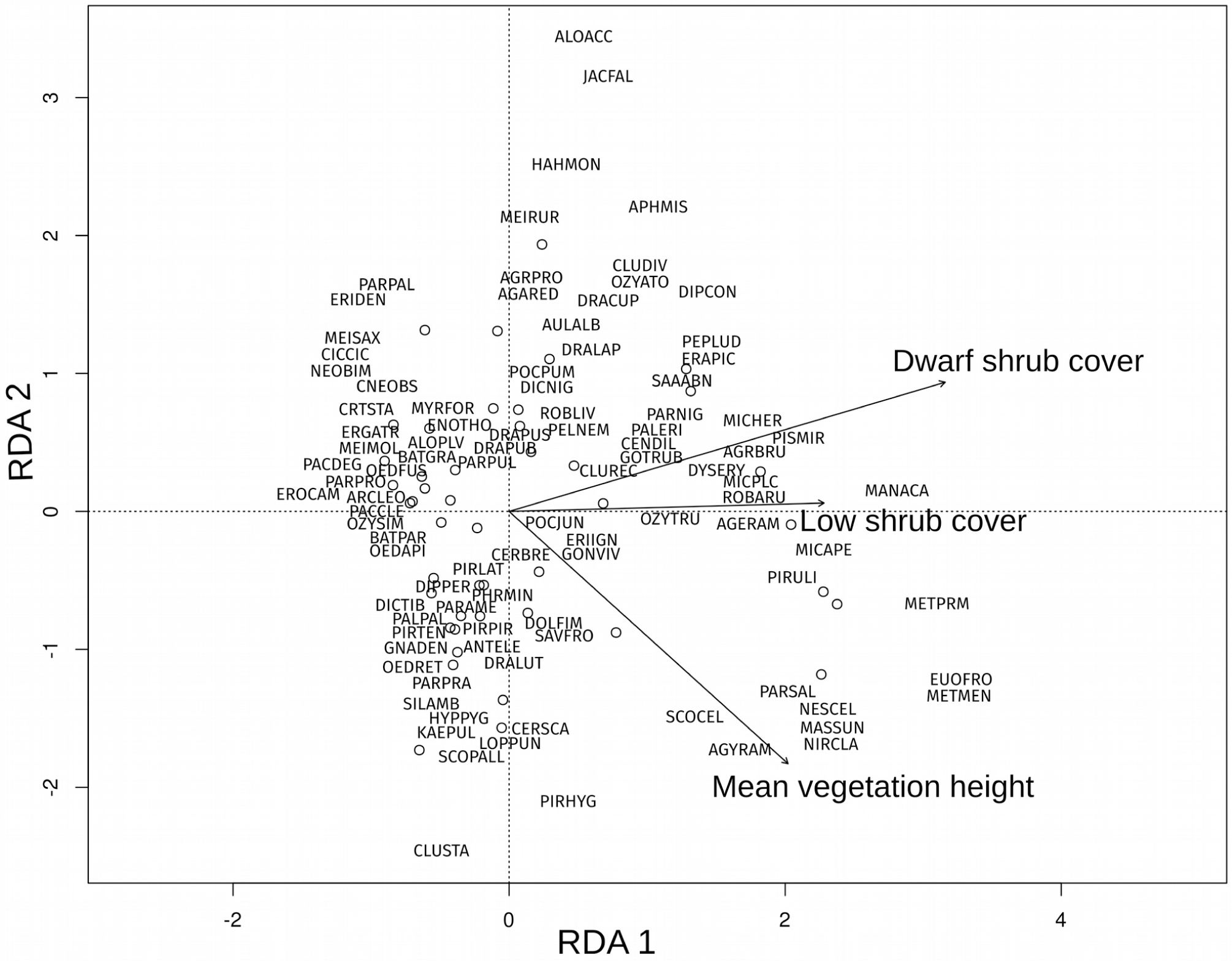
Projection of spider species and sites on the CCA axes. Circles represent sites. Species names are abbreviated as three first letters of genus and species (see species codes in Table S2).

Each set of sampling stations had a wide heterogeneity of structural forms resulting from several factors (de Fo ucault 1984; Clément and Aidoud 2009). While these vegetation forms have close spatial links, they nevertheless belong to different dynamical series mainly determined by edaphic conditions, notably water and trophic level. The diversity of past and current management practices (mostly mowing and grazing at different pressure levels) also explains the diversity of forms as encroachment or regressive vegetation stages.

Wet heaths have a progressive dynamic, ranging from dwarf shrub communities (*Ulex gallii-Erica tetralix*) to woodlands (*Molinia caerulea-Quercus robur*), with thickets (*Frangula alnus*) and birch woodlands (*Betula pubescens*) as intermediate stages. They are typically dominated by dwarf shrubs and by the Purple moor-grass (*Molinia caerulea*).

Fen meadows are characterised by low intensity management by mowing or grazing (*Caro verticillati-Molinietum caeruleae* or *Nardo strictae-Juncion squarrosi*) and in the most intensive conditions to improved grasslands (*Cynosurion cristati* or *Lolio perennis-Plantaginion majoris*). *Juncus acutiflorus* fens are dominated by rushes and medium herbs. The process of encroachment can lead to a mire characterised by megaphorbs such as *Filipendula ulmaria* and *Angelica sylvestris* with structures strongly shaped by *Molinia caerulea*. Site characteristics are summarised in Table 1 and 2. Location and type of sampling stations within each site are provided in Online Resource 1.

**Table 1:** Site (LB: fens and heathlands of Lann Bern and Magoar-Penn Vern; LG: bogs of Langazel) characteristics summarized by phytosociological association (CARJUN: *Caro verticillati-Juncetum acutiflori*; CARLAS: *Caricetum lasiocarpae* (generic); CARMOL: *Caro verticillati-Molinietum caeruleae*; CIRFES: *Cirsio dissecti-Scorzoneretum humilis;* SCOFES: *Scorzonero humilis-Festucetum asperifoliae*; SPHERI: *Sphagno compacti-Ericetum tetralicis*; ULIERI: *Ulici gallii-Ericetum tetralicis*). Mean height: mean ± SD vegetation height (cm), Max height: mean ± SD maximum vegetation height (cm). Values for Dwarf shrub, Forb, Grass, Rush and Tree are mean ± SD % cover.

| Site Association | Mean height | Max height | Dwarf shrub | Forb | Grass | Rush | Tree |
| --- | --- | --- | --- | --- | --- | --- | --- |
| LB CARJUN | 45 ± 37.19 | 130 ± 116.33 | 0.2 ± 0.4 | 0.13 ± 0.09 | 0.38 ± 0.37 | 0.00 | 0.2 ± 0.36 |
| CARMOL | 35.00 | 110.00 | 0.00 | 0.10 | 0.81 | 0.02 | 0.02 |
| CIRFES | 45.00 | 110.00 | 0.00 | 0.25 | 0.24 | 0.00 | 0.40 |
| SCOFES | 21.67 ± 10.41 | 85 ± 13.23 | 0.00 | 0.43 ± 0.37 | 0.49 ± 0.3 | 0.00 | 0.04 ± 0.04 |
| SPHERI | 65.00 | 140.00 | 0.00 | 0.42 | 0.02 | 0.00 | 0.51 |
| ULIERI | 38.85 ± 23.38 | 78.85 ± 34.89 | 0.19 ± 0.23 | 0.09 ± 0.14 | 0.48 ± 0.26 | 0.02 ± 0.05 | 0.14 ± 0.27 |
| LG CARJUN | 23.33 ± 12.11 | 64.17 ± 32.16 | 0.06 ± 0.15 | 0.18 ± 0.19 | 0.65 ± 0.23 | 0.01 ± 0.02 | 0.05 ± 0.06 |
| CARLAS | 15.00 | 25.00 | 0.00 | 0.02 | 0.66 | 0.00 | 0.06 |
| CARMOL | 53.33 ± 22.73 | 172.5 ± 171.4 | 0.36 ± 0.35 | 0.02 ± 0.04 | 0.5 ± 0.33 | 0.09 ± 0.1 | 0.02 ± 0.04 |
| CIRFES | 30 ± 5 | 58.33 ± 15.28 | 0.16 ± 0.28 | 0.05 ± 0.06 | 0.59 ± 0.23 | 0.00 | 0.09 ± 0.07 |
| SCOFES | 28.33 ± 20.21 | 63.33 ± 16.07 | 0.3 ± 0.27 | 0.13 ± 0.23 | 0.33 ± 0.13 | 0.06 ± 0.09 | 0.15 ± 0.2 |
| SPHERI | 45.00 | 55.00 | 0.36 | 0.02 | 0.55 | 0.00 | 0.01 |
| ULIERI | 40.00 | 82.5 ± 3.54 | 0.00 | 0.23 ± 0.11 | 0.55 ± 0.13 | 0.00 | 0.12 ± 0.15 |

**Table 2:** Mean ± SD Ellenberg index values by site (LB: fens and heathlands of Lann Bern and Magoar-Penn Vern; LG: bogs of Langazel) and association (CARJUN: *Caro verticillati-Juncetum acutiflori*; CARLAS: *Caricetum lasiocarpae* (generic); CARMOL: *Caro verticillati-Molinietum caeruleae*; CIRFES: *Cirsio dissecti-Scorzoneretum humilis;* SCOFES: *Scorzonero humilis-Festucetum asperifoliae*; SPHERI: *Sphagno compacti-Ericetum tetralicis*; ULIERI: *Ulici gallii-Ericetum tetralicis*). L: mean Ellenberg index value for light, F: mean Ellenberg index value for moisture, R: mean Ellenberg index value for pH, N: mean Ellenberg index value for nitrogen, S: mean Ellenberg index value for conductivity.

| Site Association | L | F | R | N | S |
| --- | --- | --- | --- | --- | --- |
| LB CARJUN | 6.83 ± 0.35 | 6.87 ± 0.45 | 3.83 ± 0.95 | 2.89 ± 0.47 | 0.01 ± 0.02 |
| CARMOL | 7.32 | 7.18 | 4.08 | 2.22 | 0.06 |
| CIRFES | 7.19 | 6.98 | 4.89 | 3.59 | 0.06 |
| SCOFES | 6.68 ± 0.28 | 6.25 ± 0.42 | 4.53 ± 0.42 | 3.45 ± 0.38 | 0.18 ± 0.03 |
| SPHERI | 6.08 | 6.29 | 4.52 | 4.04 | 0.09 |
| ULIERI | 7.13 ± 0.29 | 6.94 ± 0.63 | 3.45 ± 1.12 | 2.54 ± 0.83 | 0.04 ± 0.08 |
| LG CARJUN | 7.18 ± 0.4 | 7.17 ± 0.52 | 3.84 ± 0.89 | 2.59 ± 0.61 | 0.06 ± 0.04 |
| CARLAS | 7.43 | 7.48 | 3.79 | 2.3 | 0 |
| CARMOL | 6.95 ± 0.33 | 6.86 ± 0.69 | 3.28 ± 0.91 | 2.47 ± 0.65 | 0.06 ± 0.11 |
| CIRFES | 7.39 ± 0.21 | 7.18 ± 0.86 | 3.63 ± 1.48 | 2.56 ± 0.96 | 0.13 ± 0.17 |
| SCOFES | 7.29 ± 0.18 | 7.42 ± 0.32 | 3.51 ± 1.35 | 2.62 ± 0.95 | 0.08 ± 0.09 |
| SPHERI | 7.39 | 6.93 | 3.32 | 1.95 | 0 |
| ULIERI | 6.99 ± 0.2 | 6.75 ± 0.58 | 4.14 ± 0.76 | 2.84 ± 0.81 | 0.06 ± 0.08 |

### Sampling design

Sampling of spiders took place in May and June 2013 over 30 consecutive days using pitfall traps. Each trap was emptied every two weeks (2 sampling sessions). To compensate for this short sampling duration, we chose to increase spatial effort, as advised by Lövei & Magura (2011). Thus, forty-five plots were sampled (23 in LB site with 15 grasslands and 8 heathlands and 22 in LG site with 13 grasslands and 12 heathlands), with four traps per plot (100 mm diameter, filled with preservative solution (50% ethylene-glycol, 50% water) (Schmidt et al. 2006). Traps were placed 10 m away from each other to avoid interference between traps (Topping and Sunderland 1992).

Vegetation surveys of 25 m^2^ plots were conducted at each site to assess vegetation growth-form (dwarf shrub, low shrub, tall shrub, forb, grass, rush, sedge and tree) cover (%) following Cristea et al. (2015) and Westhoff and van der Maarel (1978). In addition, five square sub-plots of 0.25 m2 each were set within each plot. In each sub-plot, vascular plant species cover (%), mean and maximum vegetation height, and litter depth were measured to the nearest cm. Values are then averaged for each plot. Nomenclature follows Platnick (2014) and Gargominy et al. (2015) for spiders and plants respectively (Online Resources 2 and 3).

### Statistical analyses

Spider activity–density was defined as the mean number of individuals caught per trap. Spider α-diversity was estimated as mean species richness per plot. Spider β-diversity was estimated using a dissimilarity matrix (corresponding to Sørensen pair-wise dissimilarity) partitioned into its two components – species turnover (βt) and nestedness (βn) – following Baselga (2010) and using the betapart R package (Baselga and Orme 2012). Functional diversity (FD) was computed according to Villéger et al. (2008) for spider activity-density and the Gower dissimilarity matrix was computed based on seven traits: body length (Roberts, 1995), seasonality (Harvey et al. 2002), hunting technique, (Uetz et al. 1999) ballooning dispersal (Bell et al. 2005), habitat specialist/generalist (Hänggi et al 1995), preferred substrate (Buchar et al. 2002), and preferred humidity (Buchar et al. 2002). The species-by-species distance matrix could not be represented in a Euclidean space so we applied a Cailliez correction (Cailliez 1983). Analyses were performed using the FD package (Laliberté and Legendre 2010; Laliberté et al. 2014).

To investigate ground spider response to abiotic environment and vegetation characteristics, drivers of species assemblages were investigated using constrained analysis. The choice between redundancy analysis (RDA) and constrained correspondence analysis (CCA) was made according to the axis length (< 3 or > 4 respectively) of a detrended correspondence analysis (DCA) (Legendre and Gallagher 2001). Here, we chose an RDA as the first axis length was 2.88. Activity-density of all individual species were the response variables. Potential predictor variables included direct measures of vegetation structures, such as vegetation height (mean and max), litter depth, growth-form-type cover (grass, sedge, dwarf shrub, low shrub, forb, rush, and tree) and abiotic variables derived from plant communities using Ellenberg indicator values (Ellenberg et al. 1992). After correlation tests, only non significantly or poorly (i.e. R < 0.5) correlated variables were kept in the RDA (see variables in Table 1). Ellenberg indices (moisture (F), nitrogen (N), pH (R), light (L), and conductivity (S)) were derived from the phytosociological relevés, using van der Maarel’s indices instead of the abundance-dominance coefficient to minimize the weight of dominant species. Moreover, species described in the Ellenberg system as indifierent were ignored (see for e.g. Dzwonko, 2002). All other species were kept in the analyses. The Ellenberg values came from Hill et al. (2004) and were corrected for the British Isles. To obtain a more relevant description of the vegetation for the study of spider communities, i.e. illustrating the structure, a simple functional classification of plants adapted from Box (1996) was used. This type of classification results from species growth forms and medium height and includes ten types of plants: trees, tall shrubs, low shrubs, dwarf shrubs, rushes, sedges, grasses, forbs, pteridophytes and vines. The contribution of the different functional groups was calculated for each secondary plot and the mean contribution was worked out for the entire area, as for vegetation height and litter depth. Litter depth was highly correlated with vegetation height (Spearman test, R = 0.587, P < 0.001) and was not included in the analyses despite its importance for ground spiders (Uetz 1979a).

Responses of spider activity-density, α-diversity and functional diversity were tested using generalised linear models (GLMs). For activity-density we used binomial-negative distribution with a logit link. For α-diversity and functional diversity we assumed a Gaussian distribution and used a linear model. Relevant variables were selected using a stepwise model selection by AIC (Akaike 1974).

To identify the variables significantly influencing spider β-diversity, we performed a multiple regression analysis on the distance matrix of predictors (MRM) following the methods outlined in Legendre et al. (1994) using the ecodist R package (Goslee and Urban 2007). Predictors were the same as for RDA and GLMs. All statistical analyses were performed using R 3.2.3 (R Core Team 2015).

## Results

A total of 8576 spider individuals belonging to 141 species (see taxonomic list in Table S1) were caught, with a high (93.38%) ratio of adult spiders. Assemblages were dominated by two lycosids (*Pirata latitans* and *Pardosa pullata* representing 24% and 23% of adult individuals respectively).

RDA on spider assemblages was significant (F_14,30_ = 1.63, df = 14, P < 0.001) and explained 43.3% of the total variance, and the four first axes of the RDA were significant (respectively F_1,30_ = 7.42, df = 1, P < 0.001; F_1,30_ = 3.87, df = 1, P = 0.001; F_1,30_ = 2.23, df = 1, P = 0.001; F_1,30_ = 1.94, df = 1, P = 0.008). Variables explaining spider composition were mean vegetation height (F_1,30_ = 3.89, df = 1, P = 0.001), dwarf shrub cover (F_1,30_ = 5.07, df = 1, P = 0.001) and low shrub cover (F_1,30_ = 1.79, df = 1, P = 0.041) (Fig. 4).

**Figure 3.**
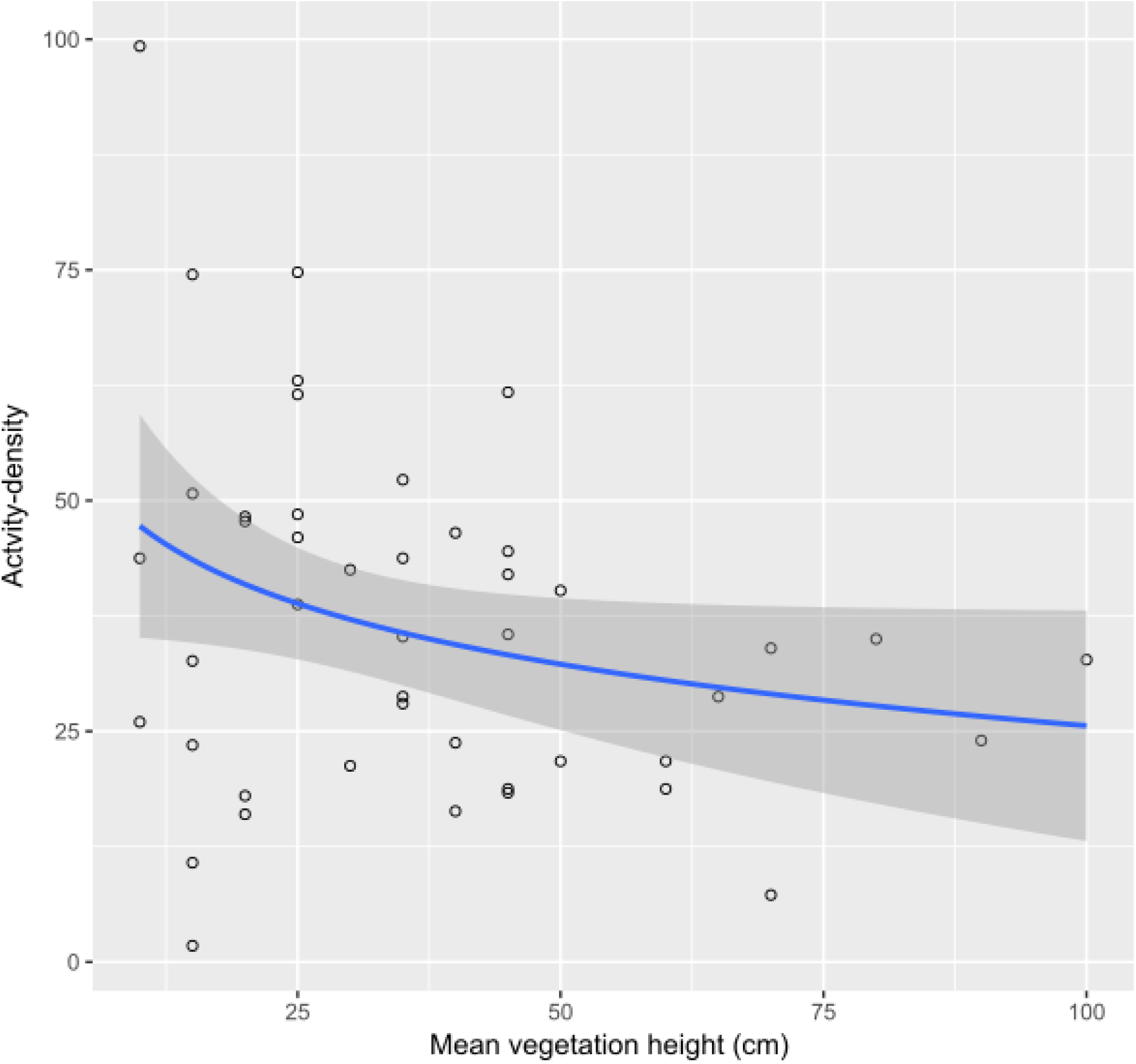
Spider activity–density as a function of mean vegetation height with log regression line and standard error.

**Figure 4.**
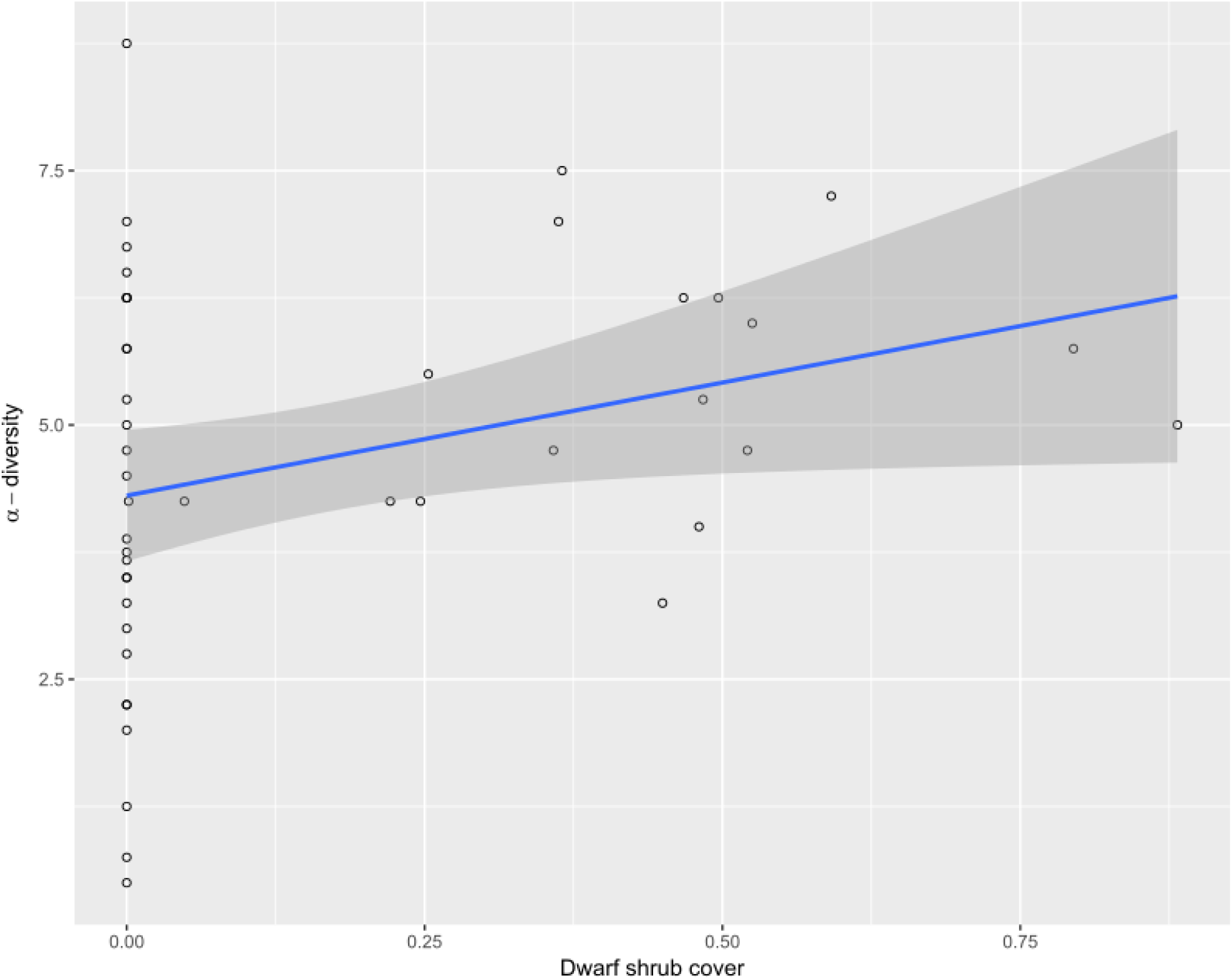
Spider α-diversity as a function of dwarf-shrub cover with log regression line and standard error.

Spider activity–density was significantly and negatively influenced by mean vegetation height and the mean Ellenberg value for moisture (Table 3). Spider α-diversity was significantly and positively influenced by dwarf shrub and forb cover (Table 3). It was also positively influenced by Ellenberg index values for light and negatively by Ellenberg index values for moisture (Table 3). Spider functional diversity was positively influenced by dwarf shrub cover and Ellenberg index values for conductivity (Table 3). MRM was significant (P = 0.007, R² = 0.13). Spider β-diversity was significantly and positively influenced by mean vegetation height and by Ellenberg index value for pH, and negatively influenced by Ellenberg index value for nitrogen (Table 3).

**Table 3.**
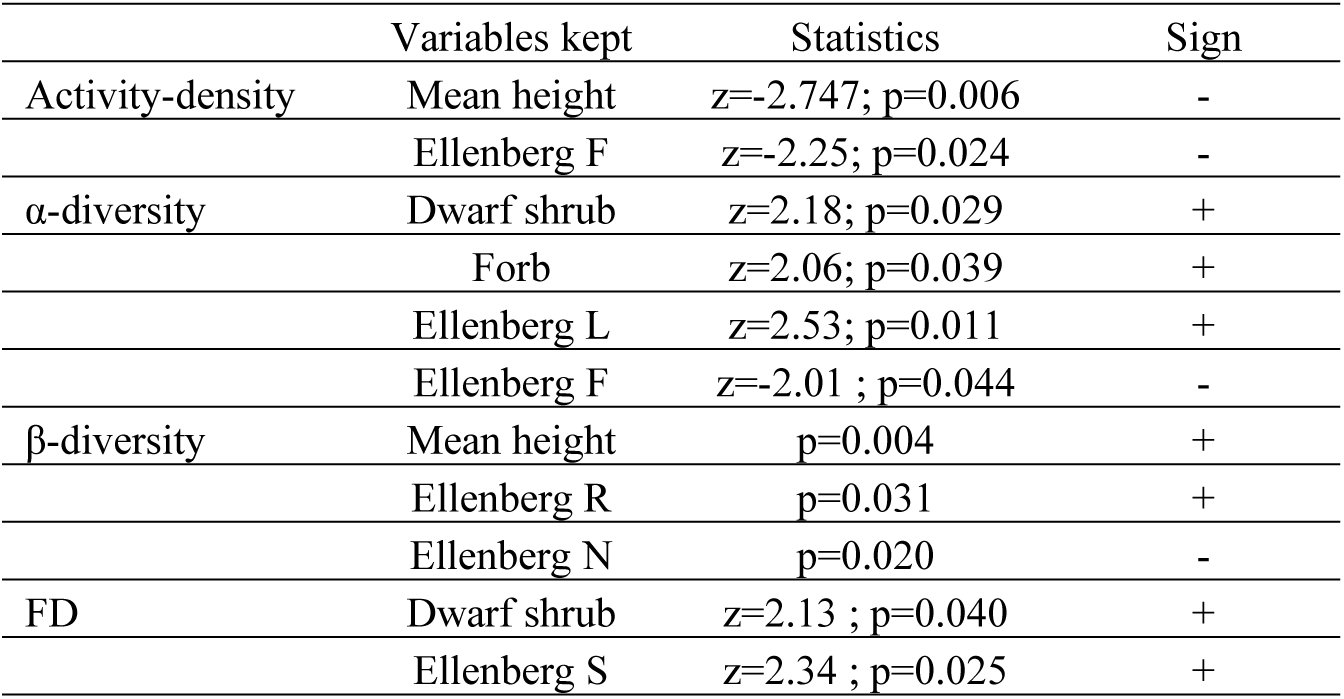
Significant explicative variables kept by the step-AIC on the results of the general linear model (GLM) on activity-density, α-diversity and functional diversity (FD) of spiders and significant explicative variables from the multiple regression on distance matrices (MRM) on spider β-diversity. Best regression models are given between brackets. (Ellenberg L: mean Ellenberg value for light; Ellenberg F: mean Ellenberg value for moisture; Ellenberg R: mean Ellenberg value for pH; Ellenberg N: mean Ellenberg value for nitrogen, S: mean Ellenberg value for conductivity; quadra: quadratic; lin: linear).

## Discussion

Our results suggest that the spider assemblages studied were strongly influenced by vegetation structure, even when considering ground-dwelling species. We found spider activity–density to be negatively influenced by vegetation height. This is in opposition with previous studies dealing with simplification of vegetation structure at both habitat and within-plant scales (see Langellotto & Denno 2004 for a meta-analysis), which found that increasing habitat complexity results in significant increases in arthropod, especially hunting and web-building spider, abundance. Predators may indeed aggregate in complex habitats because they find a higher abundance of prey, refuge from predation, better conditions for prey localisation and capture, a favourable micro-climate, and alternative resources (Langellotto & Denno 2004). Nevertheless, results for ground-dwelling spiders are usually less clear. For instance, Hurd & Fagan (1992) did not find any clear effect of succession (from herbaceous to woody communities) on spider abundance. In our dataset, vegetation height was highly and positively correlated to litter depth. Uetz (1979) found that the activity-density of Lycosidae (that were the most abundant spiders in our study) was negatively influenced by increasing litter depth. Consequently the negative relationship observed between vegetation height and spider activity-density is most likely the consequence of changes in litter characteristics rather that in vegetation structure. It is important to note that spider activity–density is dependent on individuals’ mobility. Changes in micro-climatic and structural conditions of habitats modified by vegetation height (Griffin and Yeargan 2002; Langellotto and Denno 2004) are actually known to affect the mobility of individuals, and therefore their catchability (which was already mentioned by Uetz 1976; Topping and Sunderland 1992; Lang 2000). Thus, the hypothesis that the negative link observed between spider activity–density and vegetation height is the consequence of a sampling bias cannot be rejected.

Spider activity–density was also negatively influenced by moisture, which is in accordance with previous findings (Uetz 1979b). In a literature review, Wise (1995) suggested that the abundance of spiders depends on three variables: wind, moisture, and temperature. More recently, Entling et al. (2007) and Lambeets et al. (2008) showed moisture to be an important driver of riparian spider activity-density.

Spider assemblages were best explained by variables reflecting vegetation structure, and more specifically vegetation closure and complexity (mean vegetation height, dwarf shrub cover, and low shrub cover). The importance of these variables is confirmed by the fact that we found dwarf shrub and forb cover to be positively related to α-diversity and mean vegetation height to increasing β-diversity. According to Entling et al. (2007), spider assemblages are mainly related to habitat type and depend on the shading, as well as the moisture, of habitats. Shading is obviously related to the development of shrubs and we logically found spider α-diversity to be positively influenced by light. Thus, changes in vegetation structure and shading may explain our results. Indeed, the role of habitat structure in itself has repeatedly been shown to determine the species richness of spiders (more so than the age of habitats, for example: Gibson, Hambler & Brown 1992; Hurd & Fagan 1992; Pétillon 2014). Greater vegetation complexity is likely to allow more species to co-exist by reducing inter-specific competition (Marshall and Rypstra 1999; Wise 2006). Vegetation closure is also positively linked to litter depth (e.g. Pétillon et al. 2008), which has been identified as a key variable explaining spider assemblages (Uetz 1979).

We found spider β-diversity was positively influenced by mean vegetation height and, therefore, vegetation closure. Spider β-diversity is considered higher in open habitats than in forests (Entling et al. 2007), and shading has long been identified as a major driver of β-diversity among habitats (MacArthur 1965). Nevertheless, at the beginning of the succession toward forested stages, the development of tall grasses and shrubs may increase β-diversity by providing new habitats for spiders. This is especially true for web-building species, but our results suggest this could also be true for ground-dwelling spiders. Studies dealing with small-scale drivers of arthropod β-diversity are scarce but compositional heterogeneity of spiders between samples is higher in young forest stands (Niemala et al. 1996). More recently, Sobek et al. (2009) found that habitat heterogeneity induced by tree diversity increases the β-diversity of true bugs. Spider β-diversity was also influenced positively by pH and negatively by nutrient level. Forestation of moorland is often characterised by an increase in pH, nutrient level, and conductivity (Kampichler and Platen 2004). Thus, spider β-diversity seems to be positively influenced by the abiotic consequences of shrub and tree development.

Functional diversity was also positively influenced by dwarf shrub cover and conductivity, indicating that the impact of vegetation closure is linked to modification of vegetation structure and abiotic changes induced by it. This result is not surprising as functional diversity is considered more sensitive to environmental change than taxonomic diversity (Cadotte et al. 2009; Schirmel et al. 2012; Woodcock et al. 2014). Indeed, taxonomic diversity often remains relatively stable regardless of vegetation changes (Brown et al., 2001; Schirmel et al., 2012) contrary to functional diversity (Schirmel et al., 2012). This is also in accordance with Schirmel et al. (2016), who found functional diversity to be higher in woody than herbaceous sites.

## Conclusion

Spider community assemblages appear to be driven by factors clearly linked to characteristics of vegetation and edaphic conditions such as vegetation height, shrub cover, pH, and soil richness. These parameters vary not only with vegetation dynamics but also according to vegetation management. In Europe, and especially in France, management strategies considered encroachment—and more generally natural dynamics—as negative trends for the conservation of agro-pastoral habitats leading to a loss of diversity and to the regression of their specific components. Management of wet heathlands and *Juncus acutiflorus* fens responds to this logic by increasing dwarf shrubby and herbaceous structures of vegetal communities through cutting or grazing operations. Conversely, our results suggest that allowing a controlled development of the shrub layer could have a positive impact on the diversity of certain groups such as ground-dwelling spiders. This clearly illustrate that “ecological value” of habitats and resulting management choices should be made using a pluritaxonomic approach. However, these results do not allow us to reach a conclusion on the value of a particular management strategy according to the type of habitat. Further analyses are consequently needed to test for the effect of management modalities and habitat types.

## Acknowledgements

We would like to thank Angélique Mangenot, Sandrine Alary (Association de Langazel), Pierrick Pustoc’h and Mélanie Ulliac (A.M.V. Lann Bern et Magoar) for field assistance, Cyril Courtial for help in species identification, and Farid Bensettiti and Lise Maciejewski (M.N.H.N.) for useful comments. This study was funded by “Région Bretagne”, French Ministry of Ecology, Sustainable Development and Energy and General Councils of Côtes d’Armor, Morbihan, Ille-et-Vilaine and Finistère.

